# Resistance screening and *in-silico* characterization of cloned novel RGA from multi race resistant Lentil germplasm against Fusarium wilt (*Fusarium oxysporum* f. sp. *lentis)*

**DOI:** 10.1101/2022.08.16.504179

**Authors:** K Nishmitha, Rakesh Singh, Sunil C Dubey, Jameel Akthar, Deeba Kamil

## Abstract

Fusarium wilt caused by *Fusarium oxysporum* f. sp. *lentis* (*Fol*) is the most devastating disease of lentil present worldwide and in India. Identification of multi-race fusarium wilt resistance genes and incorporation into existing cultivar will help to reduce yield loss. In the present study, a hundred lentil germplasm were screened against seven prevalent races of *Fol* and accession IC201561, EC714243 and EC718238 were identified resistant. The typical R gene codes for the nucleotide-binding site and leucine-rich repeats (NBS-LRR) at the C terminal linked to either Toll/interleukin 1-like receptor (TIR) or coiled-coil (CC) at the N terminal. In the present study degenerate primers designed from the NBS region amplifying P-loop to GLPLA motif isolated forty-five resistance gene analogues (RGA) from identified resistant accessions. The sequence alignment identified both classes of RGA, TIR and non-TIR based on the presence of Aspartate (D) and Tryptophan (W) at the end of kinase motif respectively. The phylogenetic analysis grouped RGA into six classes, LRGA1 to LRGA6 determining the diversity of RGA present in the host. Grouping of RGA identified from *Lens nigricans*, LnRGA 2, 9, 13 with I2 reveals a probable role in Fusarium resistance. The similarity index of 27.85% to 86.98% was found among RGA and 26.83% to 49.41% between known R genes, I2, Gpa2, M and L6. Active binding sites present along the conserved motifs have grouped the RGA into 13 groups. ADP/ATP being the potential ligand determines ATP binding and ATP hydrolysis activity of RGA. The isolated RGA can be used in developing marker linked to the functional R gene. Further, expression analysis and full-length gene isolation further pave path to identifying the molecular mechanism involved in resistance.

## Introduction

Lentil (*Lens culinaris* Medikus subsp. *culinaris*) is one of the most important cool-season legume food crop grown after chickpea. It is an annual, self-pollinating diploid (2n= 14) crop having a genome size of approximately 4Gb [1]. It is the oldest crop that originated in Turkey and is now cultivated in almost all parts of the world. It is majorly grown in North America, Africa, Middle East and Asia. Globally lentil is grown in an area of 6.1million ha and produce about 6.33mtons of yield. India has the largest area under cultivation of approximately 1.51 mha and placed second in production after Canada producing 1.096 million metric tons [2]. Lentil is a rich source of protein (24-26%) and fiber. It is also a source of micronutrients such as calcium, phosphorus and iron and essential amino acids such as lysine [3]. It is majorly grown as a rabi season legume crop in northern states of India such as Delhi, Uttar Pradesh, Madhya Pradesh, West Bengal, Rajasthan, Punjab and Haryana. It is also grown as a rotational rainfed crop on previous season residual moisture. It is considered as a valuable crop as it consumes minimum input yet fixing atmospheric nitrogen, enhancing soil fertility and generating means of livelihood to small scale farmer [4]. However, its yield is highly affected by biotic stress and abiotic stress. Among which Fusarium wilt caused by *Fusarium oxysporum* f. sp. *lentis* Vasudeva and Srinivasan is one of the major constrain in lentil production globally and in India. The disease causes drying of leaves and seedling death during the seedling stage and partial or complete wilting during the reproductive stage [5]. The presence of multiple races in the Indian subcontinent has made its control more challenging [6]. In India, it accounts for 100% yield loss if affected in the seedling stage [7] and 50-70% in natural conditions [8]. Resistance breeding by identification and incorporation of single or multiple race-specific resistance genes into cultivar would considerably control the disease in a geographic area. Few resistant varieties have been released previously but the potential of available germplasm has been less explored. Screening for resistance during different plant growth stages helps to identify late wilters during the reproductive stage and temporal variation of resistance [9]. Wild lentil species have been identified as a potential source of resistance that can be tailored for resistance breeding [10]. Multilocation screening of germplasm identifies germplasm resistant to multiple races present in a geographic region [11]. Conventionally, the Resistance (R) gene is isolated by transposon tagging and map-based cloning which is laborious and time-consuming. The structure of R gene is conserved across plants with a typical NBS-LRR region at the C terminal linked to Coiled-coil (CC) or Toll/interleukin 1-receptor (TIR) at N terminal end [12]. A degenerate primer designed from the conserved NBS region can be used to PCR amplify, Resistance gene analogues (RGA). Previously RGA has been cloned in lentil [13] and other legume crops such as chickpea, Fababean [14] pigeon pea [15], soybean [16] and common bean [17]. The isolated RGA mainly belonged to the TIR and non-TIR classes of NBS and serves as a marker linked to the functional R gene and helps in isolating the full-length R gene [18]. Previously no lentil germplasm has been identified resistant to multiple races of *Fol*. Isolation and characterization of potential RGA from resistant sources serve as a useful tool for full-length gene isolation. The present study was undertaken with the following objective. 1. To screen a hundred lentil accession belonging to one cultivated species and six wild species to identify the resistance source against seven races of *Fol*. 2. To isolate the RGA from resistant accession for further characterization and identify potential interacting partners.

## Materials and method

### Fungal isolates

Seven races of *Fusarium oxysporum* f. sp. *lentis* (MP-2, UP-9, RJ-8, DL-1, CH-5, UP-12 and BR-27) reported earlier [6] were used in present study. Pure culture of isolates was maintained in PDA slants and stored at 4 °C for further study.

### Plant material

One hundred accessions belonging to *Lens culinaris* subsp. *culinaris* (70), *L. c*. subsp. *tomentosus* (2), *L. c* subsp. *orientalis* (7), *L. c*. subsp. *odemensis* (5), *L. lamottei* (3), *L. nigricans* (6) and *L. ervoides* (7) were collected from ICAR-NBPGR, New Delhi. The susceptible check (L-9-12) and resistant check (PL639) were collected from Division of Genetics, ICAR-IARI, New Delhi.

### Screening and evaluation for resistance

The experiment was carried out in ICAR-NBPGR, New Delhi during 2020-21 and 2021-22 as previously described method by Bayaa and Erikson [9]. The seven races of *Fusarium oxysporum* f. sp. *lentis* were grown in double autoclaved sorghum seeds for fifteen days. Fifteen grams of inoculum were mixed in pots containing 2kg of sterilized soil. Surface sterilized seeds of a hundred accession were sown in the pots and eight plants/ pot were maintained at 24°C/22°C with their respective control (untreated). The accessions IC73121 against BR-27, IC95658 against UP-9, IC361467 against CG-5, IC384447 and IC53238 against UP-12, EC718234 against MP-2, CG-5, UP-12 and EC718330 against DL-1and UP-12 showed poor germination after repeated sowing and were not included carried further. The performance of disease pressure was compared with resistant and susceptible checks. Disease incidence was recorded every week until the pod filling stage and a scale of 1-9 was used [19] to identify resistant accession for further RGA isolation and characterization.

### Genomic DNA isolation and PCR amplification

Genomic DNA of three accessions, IC201561, EC714243 and EC718238 showing resistance to the majority of races were isolated using the modified CTAB method [20]. Quality and quantity of DNA was checked in 0.8% agarose gel electrophoresis and nanodrop and stored at -20°C. Previously designed degenerate primer from the conserved NBS region of R gene amplifying P-loop to GLPLA motif was used in the present study [13] (S1 Table). PCR reaction of 25µl was carried out in 0.2ml PCR tubes containing 10x Taq buffer (Thermo Fisher), 0.5µl of 10mM dNTP’s, 10pmol of each degenerate primer, 1U of Taq polymerase (Thermo Fisher) and 100ng of template. PCR amplification was carried out with specific conditions of initial denaturing at 95°C for 5 min, followed by 36 cycles of denaturation at 95°C for 1min, annealing at 45°C for 1 min, elongation at 72°C for 1 min, final elongation for 5 min followed and cooling at 4°C in a thermal cycler (Bio-rad). The amplified products were visualized on 1.2 % agarose gel electrophoresis. PCR product corresponding to 510 bp was eluted and purified using QIAquick Gel Extraction kit (Qiagen, Hilden, Germany).

### Cloning and sequencing

Purified PCR product was ligated to pGEMT easy vector (Promega, Madison, Wis.) and cloned to E. coli JM109 according to manufactures protocol. About fifty positive colonies from each transformation were screened using colony PCR. Clones producing a band of ^≈^510bp were further proceeded for plasmid isolation and EcoR1 (Thermo Fisher) restriction digestion. Plasmids were isolated using Wizard Plus Plasmid Minipreparation Kit (Promega) and sequenced by outsourcing.

### In-sillico characterization

Obtained sequences were trimmed for vector contamination and a similarity search was performed using BLAST algorithm in GenBank database. The amino acid sequences were deduced using Expasy. Multiple alignment of obtained amino acids was carried out using CLUSTALX in BioEdit software. Phylogenetic tree was constructed by Neighbor-joining method [21] with Poisson correction in MEGAX software along with NBS region of known R gene, N (U15605), L6 (U27081), M (U73916), RPP5 (AAFO8790.1), RPP4 (AAM18462.1), RPP1 (AT3444670), RPS4 (CAB50708.1), Mla (AAG37356), Pi-ta (ACY24970.1), Pi36 (ADF29629.1), Pib (BAA76282.2), I2 (AF004878), RPP13 (AAF42831), RPM1 (AQ39214), Prf (U65391), Gpa2 (AF195939), RPP8 (AAC78631.1.), FOM-2 (AY583855.1) [22]. The confidence value was checked by bootstrapping 1,000 replicates from the original data. The percent sequence similarity, between representative RGA from each class and among known R gene, L6, M, I2 and Gpa2 were determined by DNAMAN 8 software using Needleman and Wunsch (Global model) and Pam matrix score. Multiple Expectation maximizations for Motif Elicitation (MEME) was used for motif identification and characterization compared with the known R gene [23]. The active binding site of RGA and its potential ligand were determined using web based I-TASSER software [24-25]. The relationship between ABS was determined by constructing a phylogenetic tree using Maximum likelihood in MEGA X. Secondary structure of RGA with percent alpha helix, beta strands and presence of transmembrane helix was determined by Pyre 2 Software [26]. The tertiary structure of RGA was determined using I-TASSER software based on C-score, TM score and RMSD (root mean square deviation) best-predicted tertiary structure was selected.

## Result

### Screening of germplasm against *Fol* races

One hundred accessions of lentil were screened against seven races of Fusarium wilt from seedling to pod filling stage at 7 days interval for two cropping seasons, 2020-2021 and 2021-2022 and were graded into 5 classes with scale of 1-9 based on disease incidence (DI). Accessions showed varying degrees of resistance to races of Fol (S2 and S3 Tables). The number of accessions exhibiting high resistance (HR) responses were 24 (*L. culinaris* subsp. *culinaris*), 26 (*L. c*. subsp. *tomentosus*), 39 (*L. c* subsp. *orientalis*), 27 (*L. culinaris* sub sp. *odemensis*), 17 (*L. lamottei*), 39 (*L. nigricans*) and 26 (*L. ervoides*) in 2020-21 and 21 (*L. culinaris* subsp. *culinaris*), 24 (*L. c*. subsp. *tomentosus)*, 38 (*L. c* subsp. *orientalis*), 26 (*L. culinaris* sub sp. *odemensis)*, 17 (*L. lamottei*), 38 (*L. nigricans*) and 25 (*L. ervoides*) in 2021-22 against race 1 to 7 respectively. Wild species, *L. culinaris* sub sp. *odemensis* (EC714243) showed resistance to all the races of *Fol* in both seasons. Accession belonging to *L. culinaris* subsp. *culinaris* (IC201693 and IC241532) were found susceptible to all the races of *Fol*.

Accessions of *L. culinaris* subsp. *culinaris* and *L. culinaris* sub sp. *orientalis* showed the most diverse reaction with scale of 1-9 and mean disease incidence (DI) of 4.85-7.20±0.29-0.32 and 3.00-6.67±1.2-1.9 respectively in 2020 and 4.88-7.22±0.28-0.36 and 3.00-6.67±1.2-1.9 in 2021 to all the races of Fol. All the accessions belonging *L. c* sub sp. *tomentosa* were highly resistant to Race 3 (RJ-8) and 7 (BR-27) with mean DI of 1.00±0.0 during both seasons. All the accession of *L. lamottei* were highly resistant to Race 3 (RJ-8) with mean DI 1.00±0.0 during 2020 and 2021. Contrastingly, it showed moderate susceptible to susceptible reaction with mean DI of 7.67±0.6 to race 5 (CG-5) and race 7 (BR-27) (Tables 1 and 2). Variation in the mean DI between races might be associated with the virulence of race and differential interplay between host and race. The CV value of wild species showed large variation due to less sample size and varied disease reactions within species (Figs 1 and 2). Few accessions showed variation in disease incidence in two seasons might be due to varied environmental conditions such as temperature.

**Table 1.**
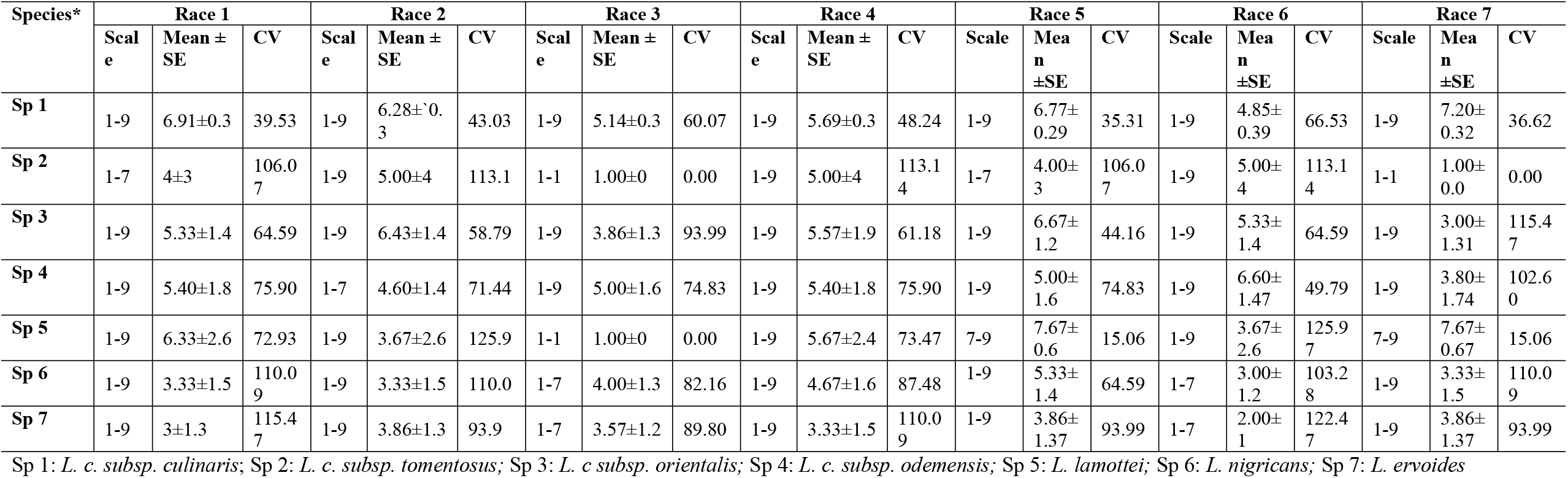
Range of variation observed in lentil germplasm against seven races of *Fusarium oxysporum* f. sp. *lentis* in the year 2020-21.

| Species* | Race 1 |  |  | Race 2 |  |  | Race 3 |  |  | Race 4 |  |  | Race 5 |  |  | Race 6 |  |  | Race 7 |  |  |
| --- | --- | --- | --- | --- | --- | --- | --- | --- | --- | --- | --- | --- | --- | --- | --- | --- | --- | --- | --- | --- | --- |
| | Scale | Mean ± SE | CV | Scale | Mean ± SE | CV | Scale | Mean ± SE | CV | Scale | Mean ± SE | CV | Scale | Mean ± SE | CV | Scale | Mean ± SE | CV | Scale | Mean ± SE | CV |
| Sp 1 | 1-9 | 6.91±0.3 | 39.53 | 1-9 | 6.28±0.3 | 43.03 | 1-9 | 5.14±0.3 | 60.07 | 1-9 | 5.69±0.3 | 48.24 | 1-9 | 6.77±0.29 | 35.31 | 1-9 | 4.85±0.39 | 66.53 | 1-9 | 7.20±0.32 | 36.62 |
| Sp 2 | 1-7 | 4±3 | 106.07 | 1-9 | 5.00±4 | 113.1 | 1-1 | 1.00±0 | 0.00 | 1-9 | 5.00±4 | 113.14 | 1-7 | 4.00±3 | 106.07 | 1-9 | 5.00±4 | 113.14 | 1-1 | 1.00±0.0 | 0.00 |
| Sp 3 | 1-9 | 5.33±1.4 | 64.59 | 1-9 | 6.43±1.4 | 58.79 | 1-9 | 3.86±1.3 | 93.99 | 1-9 | 5.57±1.9 | 61.18 | 1-9 | 6.67±1.2 | 44.16 | 1-9 | 5.33±1.4 | 64.59 | 1-9 | 3.00±1.31 | 115.47 |
| Sp 4 | 1-9 | 5.40±1.8 | 75.90 | 1-7 | 4.60±1.4 | 71.44 | 1-9 | 5.00±1.6 | 74.83 | 1-9 | 5.40±1.8 | 75.90 | 1-9 | 5.00±1.6 | 74.83 | 1-9 | 6.60±1.47 | 49.79 | 1-9 | 3.80±1.74 | 102.60 |
| Sp 5 | 1-9 | 6.33±2.6 | 72.93 | 1-9 | 3.67±2.6 | 125.9 | 1-1 | 1.00±0 | 0.00 | 1-9 | 5.67±2.4 | 73.47 | 7-9 | 7.67±0.6 | 15.06 | 1-9 | 3.67±2.6 | 125.97 | 7-9 | 7.67±0.67 | 15.06 |
| Sp 6 | 1-9 | 3.33±1.5 | 110.09 | 1-9 | 3.33±1.5 | 110.0 | 1-7 | 4.00±1.3 | 82.16 | 1-9 | 4.67±1.6 | 87.48 | 1-9 | 5.33±1.4 | 64.59 | 1-7 | 3.00±1.2 | 103.28 | 1-9 | 3.33±1.5 | 110.09 |
| Sp 7 | 1-9 | 3±1.3 | 115.47 | 1-9 | 3.86±1.3 | 93.9 | 1-7 | 3.57±1.2 | 89.80 | 1-9 | 3.33±1.5 | 110.09 | 1-9 | 3.86±1.37 | 93.99 | 1-7 | 2.00±1 | 122.47 | 1-9 | 3.86±1.37 | 93.99 |

Sp 1: *L. c. subsp. culinaris*; Sp 2: *L. c. subsp. tomentosus*; Sp 3: *L. c. subsp. orientalis*; Sp 4: *L. c. subsp. odemensis*; Sp 5: *L. lamottei*; Sp 6: *L. nigricans*; Sp 7: *L. ervoides*

**Table 2.**
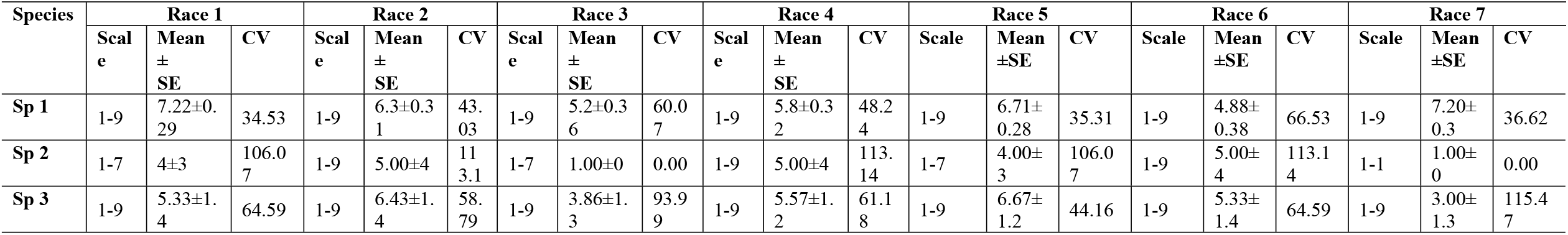
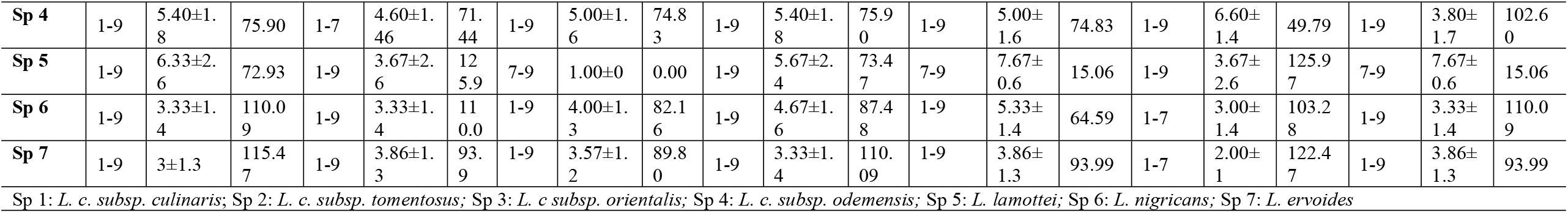
Range of variation observed in lentil germplasm against seven races of *Fusarium oxysporum* f. sp. *lentis* in the year 2021-22.

| Species | Race 1 |  |  | Race 2 |  |  | Race 3 |  |  | Race 4 |  |  | Race 5 |  |  | Race 6 |  |  | Race 7 |  |  |
| --- | --- | --- | --- | --- | --- | --- | --- | --- | --- | --- | --- | --- | --- | --- | --- | --- | --- | --- | --- | --- | --- |
| | Scale | Mean ± SE | CV | Scale | Mean ± SE | CV | Scale | Mean ± SE | CV | Scale | Mean ± SE | CV | Scale | Mean ± SE | CV | Scale | Mean ± SE | CV | Scale | Mean ± SE | CV |
| Sp 1 | 1-9 | 7.22±0.29 | 34.53 | 1-9 | 6.3±0.31 | 43.03 | 1-9 | 5.2±0.36 | 60.07 | 1-9 | 5.8±0.32 | 48.24 | 1-9 | 6.71±0.28 | 35.31 | 1-9 | 4.88±0.38 | 66.53 | 1-9 | 7.20±0.3 | 36.62 |
| Sp 2 | 1-7 | 4±3 | 106.07 | 1-9 | 5.00±4 | 113.1 | 1-7 | 1.00±0 | 0.00 | 1-9 | 5.00±4 | 113.14 | 1-7 | 4.00±3 | 106.07 | 1-9 | 5.00±4 | 113.14 | 1-1 | 1.00±0 | 0.00 |
| Sp 3 | 1-9 | 5.33±1.4 | 64.59 | 1-9 | 6.43±1.4 | 58.79 | 1-9 | 3.86±1.3 | 93.99 | 1-9 | 5.57±1.2 | 61.18 | 1-9 | 6.67±1.2 | 44.16 | 1-9 | 5.33±1.4 | 64.59 | 1-9 | 3.00±1.3 | 115.47 |
| **Sp 4** | 1-9 | 5.40±1.8 | 75.90 | 1-7 | 4.60±1.46 | 71.44 | 1-9 | 5.00±1.6 | 74.83 | 1-9 | 5.40±1.8 | 75.90 | 1-9 | 5.00±1.6 | 74.83 | 1-9 | 6.60±1.4 | 49.79 | 1-9 | 3.80±1.7 | 102.60 |
| **Sp 5** | 1-9 | 6.33±2.6 | 72.93 | 1-9 | 3.67±2.6 | 125.97 | 7-9 | 1.00±0 | 0.00 | 1-9 | 5.67±2.4 | 73.47 | 7-9 | 7.67±0.6 | 15.06 | 1-9 | 3.67±2.6 | 125.97 | 7-9 | 7.67±0.6 | 15.06 |
| **Sp 6** | 1-9 | 3.33±1.4 | 110.09 | 1-9 | 3.33±1.4 | 110.09 | 1-9 | 4.00±1.3 | 82.16 | 1-9 | 4.67±1.6 | 87.48 | 1-9 | 5.33±1.4 | 64.59 | 1-7 | 3.00±1.4 | 103.28 | 1-9 | 3.33±1.4 | 110.09 |
| **Sp 7** | 1-9 | 3±1.3 | 115.47 | 1-9 | 3.86±1.3 | 93.9 | 1-9 | 3.57±1.2 | 89.80 | 1-9 | 3.33±1.4 | 110.09 | 1-9 | 3.86±1.3 | 93.99 | 1-7 | 2.00±1 | 122.47 | 1-9 | 3.86±1.3 | 93.99 |

Sp 1: *L. c. subsp. culinaris*; Sp 2: *L. c. subsp. tomentosus*; Sp 3: *L. c. subsp. orientalis*; Sp 4: *L. c. subsp. odemensis*; Sp 5: *L. lamottei*; Sp 6: *L. nigricans*; Sp 7: *L. ervoides*

**Fig 1.**
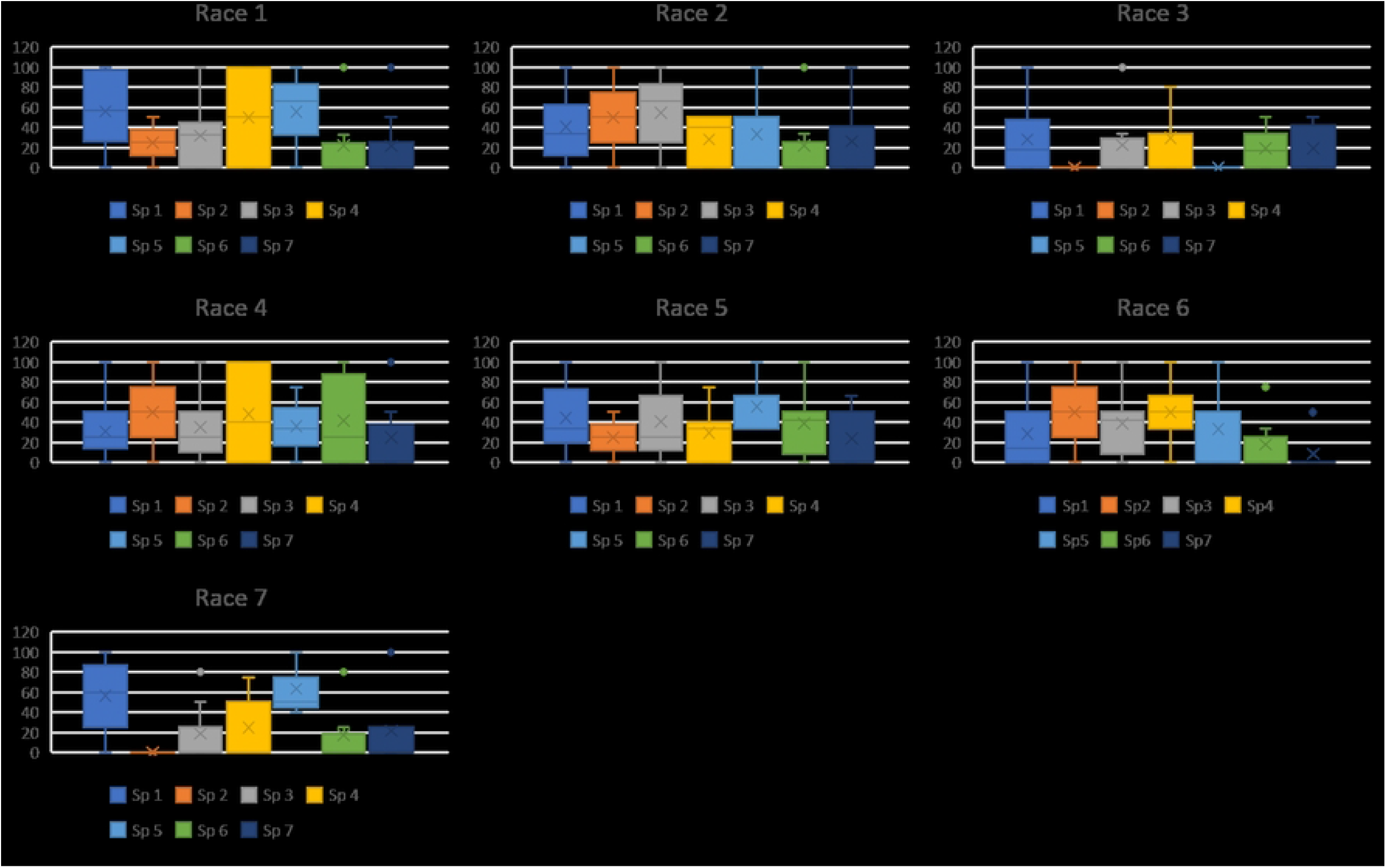
Box plot depicting disease incidence of each species of lentil against each race of *Fusarium oxysporum* f. sp. *lentis* screened during 2020-2021. Sp1: *Lens culinaris* sub sp. *culinaris*; Sp2: *L. c*. subsp. *tomentosus;* Sp3: *L. c* sub sp. *orientalis*; Sp 4: *L. c*. sub sp. *odemensis*; Sp 5: *L. lamottei*: Sp 6: *L. nigricans*; Sp 7: *L. ervoides*; Race1: MP-2; Race 2: UP-9; Race 3: RJ-8; Race 4: DL-1; Race 5: CG-5; Race 6: UP-12; Race 7: BR-27.

**Fig 2.**
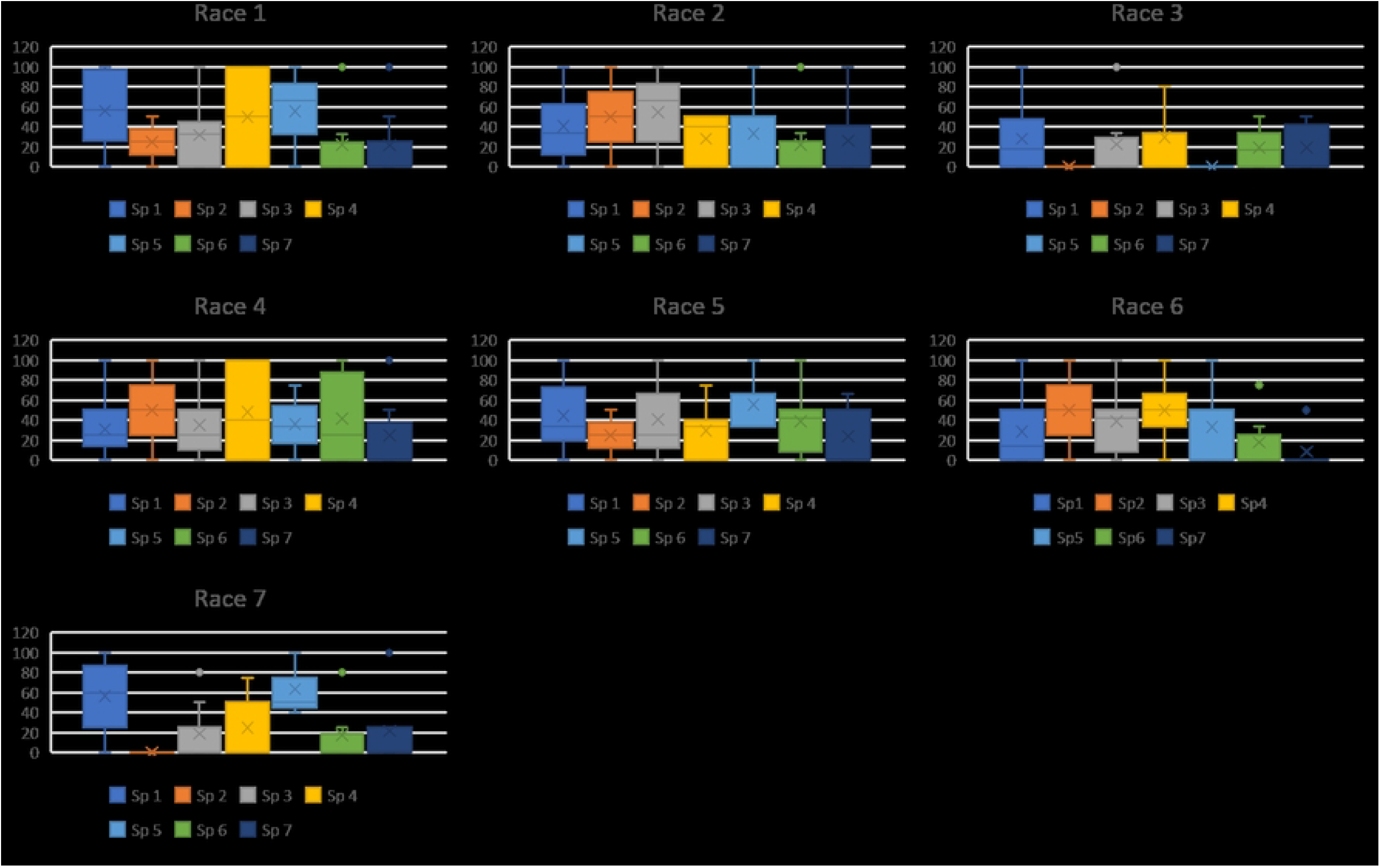
Box plot depicting disease incidence of each species of lentil against each race of *Fusarium oxysporum* f. sp. *lentis* screened during 2021-2022. Sp1: *Lens culinaris* sub sp. *culinaris*; Sp2: *L. c*. subsp. *tomentosus;* Sp 3: *L. c* sub sp. *orientalis*; Sp 4: *L. c*. sub sp. *odemensis*; Sp 5: *L. lamottei*: Sp 6: *L. nigricans*; Sp 7: *L. ervoides*; Race1: MP-2; Race 2: UP-9; Race 3: RJ-8; Race 4: DL-1; Race 5: CG-5; Race 6: UP-12; Race 7: BR-27.

### Amplification and cloning of RGA

After screening the accessions with seven races of *Fol* the accessions IC201561 (*L. culinaris*. subsp. *culinaris*), EC714243 (*L. c*. subsp. *odemensis*.) and EC718238 (*L. nigricans*) were showing resistance response to *Fol* and were used for RGA isolation and characterization (S1 Figure). The degenerate primers were designed based on conserved region of NBS amplifying P-loop to GLPLA region (S1 Table) and used for PCR amplification of genomic DNA of selected resistant accession, IC201561, EC714243 and EC718238. Amplicon size of 510 bp was eluted and cloned to pGEMT vector and E. coli JM109 cells (Fig 3). Fifty positive clones from each accession were selected for colony PCR and clones producing ^≈^510 bp further proceeded for plasmid isolation. Plasmids were isolated and restrict digested using Eco R1 to confirm insert and sequenced (S2, S3 and S4 Figures). Out of ninety sequences, 45 sequences showed high similarity to known R gene and were deposited in NCBI database (Table 3). These sequences were translated using Expasy and the sequences showed similarity ranging from 76-91% similarity with known R gene, RUN1 and RRP13 of *Medicago truncatula* and N of *Trifolium partense* and 84-99% similarity with previously isolated lentil, pea and French bean RGA (Table 3).

**Table 3.** Accession number obtained for isolated lentil RGA and result of similarity search between LRGA and known R gene from other plant species using BLASTX.

| Lentil RGA | Accession number | Similarity with other R gene | Maximum identity | E-value |
| --- | --- | --- | --- | --- |
| LcRGA1 | ON367522 | RUN1 disease resistance protein *Medicago truncatula* | 83 | 7e-125 |
| LcRGA2 | ON420341 | RUN1 disease resistance protein *M. truncatula* | 99 | 5e-186 |
| LcRGA3 | ON367523 | RUN1 disease resistance protein *M. truncatula* | 83 | 2e-124 |
| LcRGA4 | ON420342 | RUN1 disease resistance protein *M. truncatula* | 86 | 5e-146 |
| LcRGA5 | ON420343 | RUN1 disease resistance protein *M. truncatula* | 88 | 2e-164 |
| LcRGA6 | ON367524 | RUN1 disease resistance protein *M. truncatula* | 83 | 2e-124 |
| LcRGA7 | ON367525 | RUN1 disease resistance protein *M. truncatula* | 83 | 2e-124 |
| LcRGA8 | ON367526 | RUN1 disease resistance protein *M. truncatula* | 83 | 2e-124 |
| LcRGA9 | ON367527 | RUN1 disease resistance protein *M. truncatula* | 83 | 2e-124 |
| LcRGA10 | ON381744 | N disease resistance protein of *Trifolium partense* | 78 | 3e-79 |
| LcRGA11 | ON381743 | RUN1 disease resistance protein *M. truncatula* | 76 | 3e-64 |
| LcRGA12 | ON367528 | RUN1 disease resistance protein *M. truncatula* | 83 | 5e-126 |
| LcRGA13 | ON367529 | RUN1 disease resistance protein *M. truncatula* | 83 | 1e-122 |
| LcRGA14 | ON420344 | N disease resistance protein of *T. partense* | 88 | 5e-166 |
| LcRGA15 | ON367530 | RUN1 disease resistance protein *M. truncatula* | 82 | 2e-119 |
| LoRGA1 | ON381745 | RUN1 disease resistance protein *M. truncatula* | 90 | 0 |
| LoRGA2 | ON399215 | RUN1 disease resistance protein *M. truncatula* | 90 | 0 |
| **LoRGA3** | ON399211 | RUN1 disease resistance protein *M. truncatula* | 91 | 0 |
| **LoRGA4** | ON399212 | RUN1 disease resistance protein *M. truncatula* | 90 | 0 |
| **LoRGA5** | ON399213 | RUN1 disease resistance protein *M. truncatula* | 90 | 0 |
| **LoRGA6** | ON399214 | RUN1 disease resistance protein *M. truncatula* | 90 | 0 |
| **LoRGA7** | ON409891 | RUN1 disease resistance protein *M. truncatula* | 90 | 6e-167 |
| **LoRGA8** | ON409892 | RUN1 disease resistance protein *M. truncatula* | 90 | 2e-167 |
| **LoRGA9** | ON409893 | RUN1 disease resistance protein *M. truncatula* | 90 | 0 |
| **LoRGA10** | ON409894 | RUN1 disease resistance protein *M. truncatula* | 90 | 2e-167 |
| **LoRGA11** | ON409895 | RUN1 disease resistance protein *M. truncatula* | 90 | 2e-167 |
| **LoRGA12** | ON409896 | RUN1 disease resistance protein *M. truncatula* | 90 | 0 |
| **LoRGA13** | ON409897 | RUN1 disease resistance protein *M. truncatula* | 90 | 0 |
| **LoRGA14** | ON409898 | RUN1 disease resistance protein *M. truncatula* | 90 | 0 |
| **LoRGA15** | ON409899 | RUN1 disease resistance protein *M. truncatula* | 89 | 3e-160 |
| **LnRGA1** | ON420345 | N disease resistance protein of *T. partense* | 89 | 2e-169 |
| **LnRGA2** | ON420346 | RPP13 disease resistance protein *M. truncatula* | 81 | 2e-113 |
| **LnRGA3** | ON420347 | RUN1 disease resistance protein *M. truncatula* | 90 | 0 |
| **LnRGA4** | ON420348 | RUN1 disease resistance protein *M. truncatula* | 90 | 2e-167 |
| **LnRGA5** | ON420349 | RUN1 disease resistance protein *M. truncatula* | 88 | 5e-166 |
| **LnRGA6** | ON454553 | RUN1 disease resistance protein *M. truncatula* | 90 | 5e-165 |
| **LnRGA7** | ON454554 | N disease resistance protein of *T. partense* | 88 | 5e-166 |
| **LnRGA8** | ON454555 | RUN1 disease resistance protein *M. truncatula* | 90 | 5e-165 |
| **LnRGA9** | ON454556 | RPP13 disease resistance protein *M. truncatula* | 81 | 2e-109 |
| **LnRGA10** | ON454557 | RUN1 disease resistance protein *M. truncatula* | 80 | 3e-105 |
| **LnRGA11** | ON454558 | RUN1 disease resistance protein *M. truncatula* | 90 | 0 |
| **LnRGA12** | ON454559 | RUN1 disease resistance protein *M. truncatula* | 90 | 2e-166 |
| **LnRGA13** | ON454562 | RUN1 disease resistance protein *M. truncatula* | 88 | 1e-166 |
| **LnRGA14** | ON454560 | RUN1 disease resistance protein *M. truncatula* | 90 | 2e-167 |
| **LnRGA15** | ON454561 | N disease resistance protein of *T. partense* | 89 | 2e-169 |

**Fig 3.**
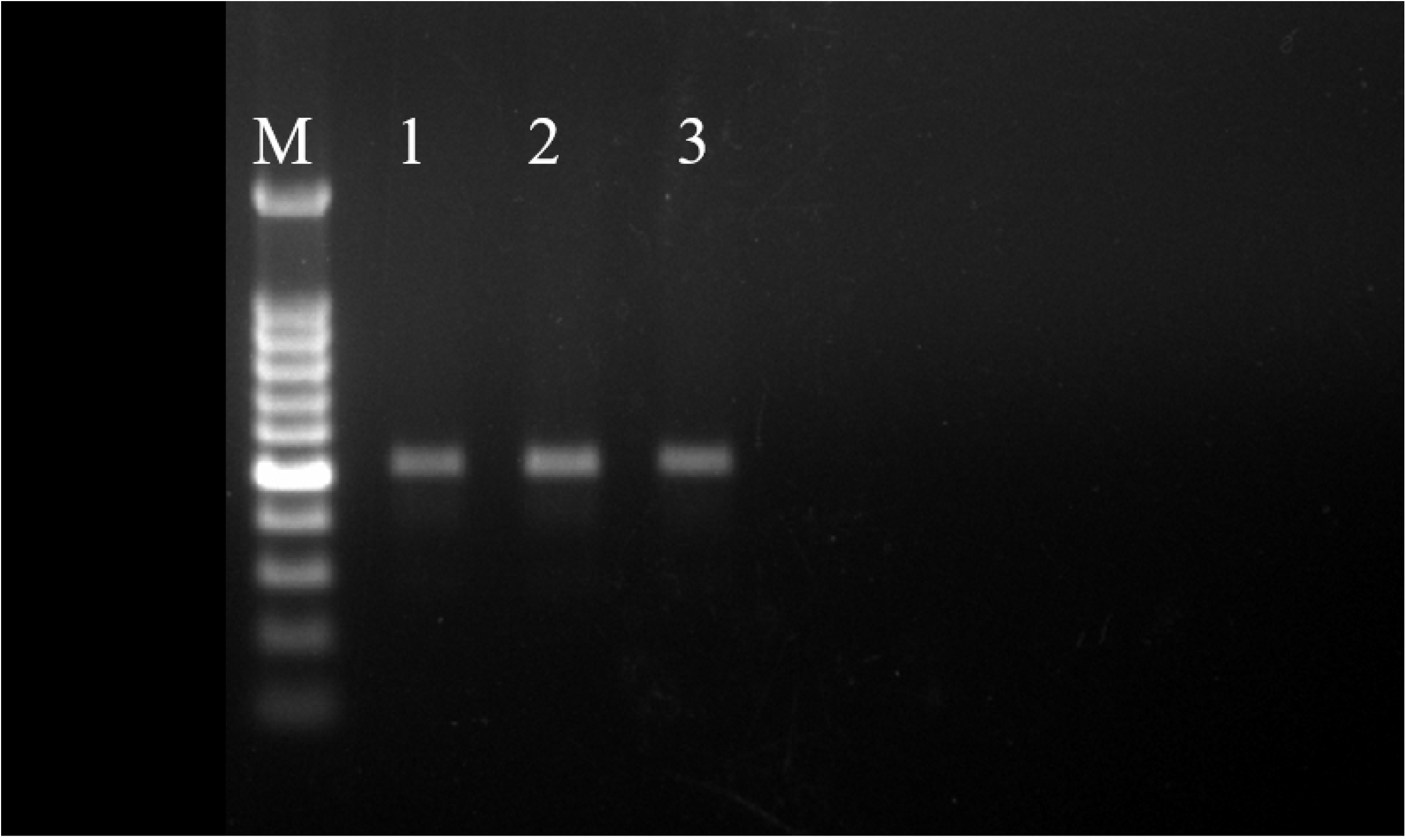
PCR amplification product generated using LRGAF and LRGAR degenerate primer on genomic DNA of lentil resistant accession. Lane 1 IC201561 (L65), Lane 2 EC714243 (L83) and Lane 3 EC718238 (L90). L represents 100 bp ladder.

### Multiple sequence alignment and phylogenetic analysis of RGAs

Multiple sequence alignment of the deduced amino acid sequence of RGA and known R gene, N, L6, M, I2 and Gpa2 revealed the presence of conserved motifs such as P-loop, RBNS-A, Kinsae2, kinase 3, RBNS-C and GLPLA (Fig 4).

**Fig 4.**
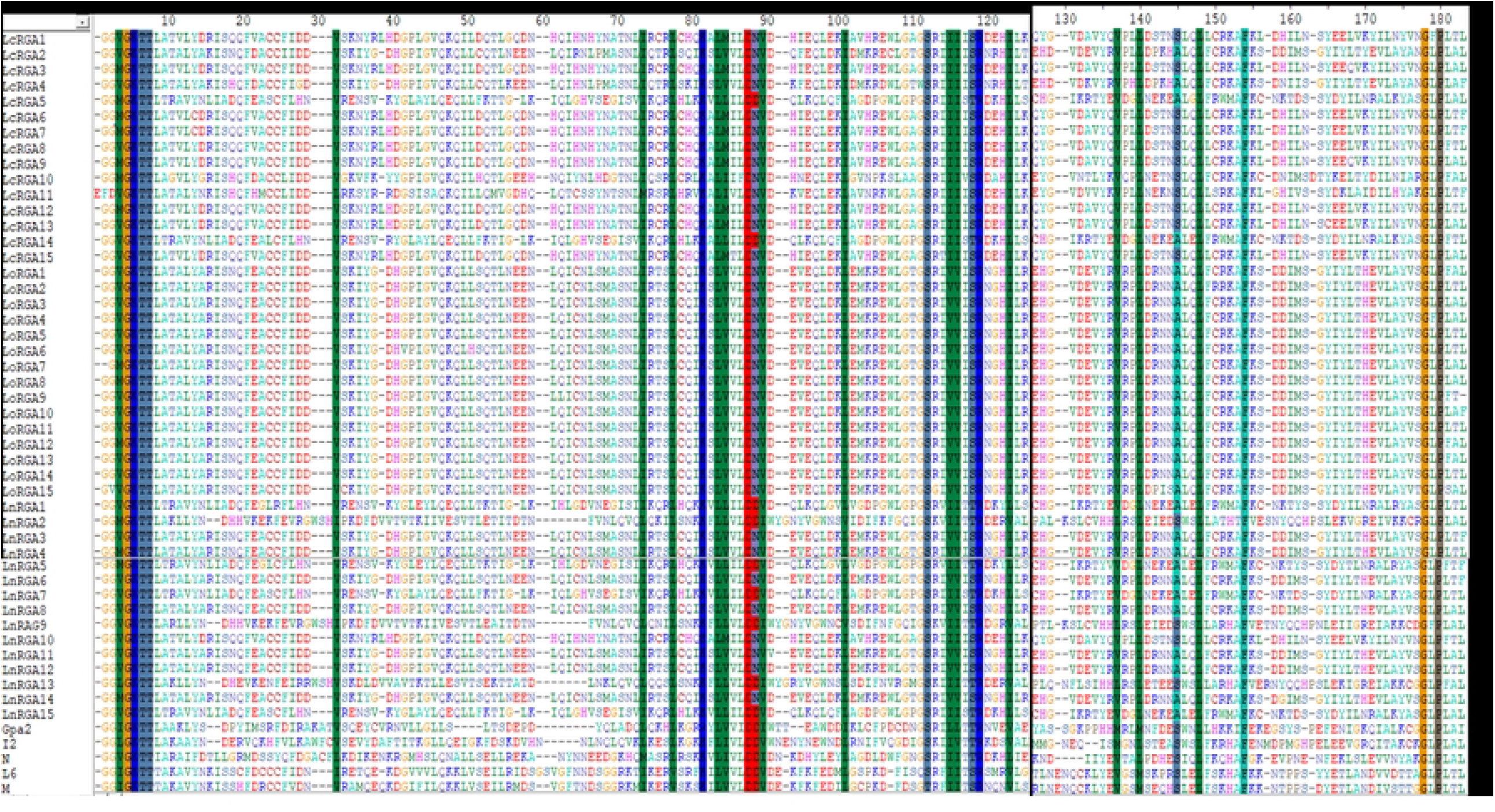
Multiple sequence alignment between P-loop and GLPLA of 45 lentil RGAs with NBS region of known R gene Gpa2, I2, N, L6 and M. The conserved motifs are indicated above the alignment. The number of amino acids is indicated above the alignment. The gaps to optimize the alignment are designated by dash (-). The alignment was constructed using CLUSTA W of Bioedit.

The phylogenetic tree was constructed using the Neighbour-joining method to determine the relationship between obtained RGA and the known R gene. The resulting tree gave rise to two branches, TIR-NBS-LRR and non-TIR-NBS-LRR. All the lentil RGA were grouped among TIR branch except LnRGA2, LnRGA9 and LnRGA13 were grouped in non-TIR branch. All lentil TIR-RGAs, R gene-L and M clustered separately from RPP4, RPP5, RPP1, N and RPS4. TIR-RGA were further classified into five classes LRGA1 to LRGA5 comprising of 22, 2, 1, 11 and 6 lentil RGA in each class. All the LoRGAs isolated from *L. c*. subsp. *odemensis* clustered together in LRGA1 reflecting sequence homology among them. RGA isolated from cultivated species and wild species were grouped in LRGA 4 and LRGA 5 revealing its conserved nature. Clustering of LnRGA3, 9 and 13 to class LRGA6 along with Fusarium R gene, I2 reveals its sequence homology and potential role in fusarium wilt resistance. Grouping of isolated RGAs to TIR and non-TIR classes reflects the diversity of RGA present in the genus (Fig 5).

**Fig 5.**
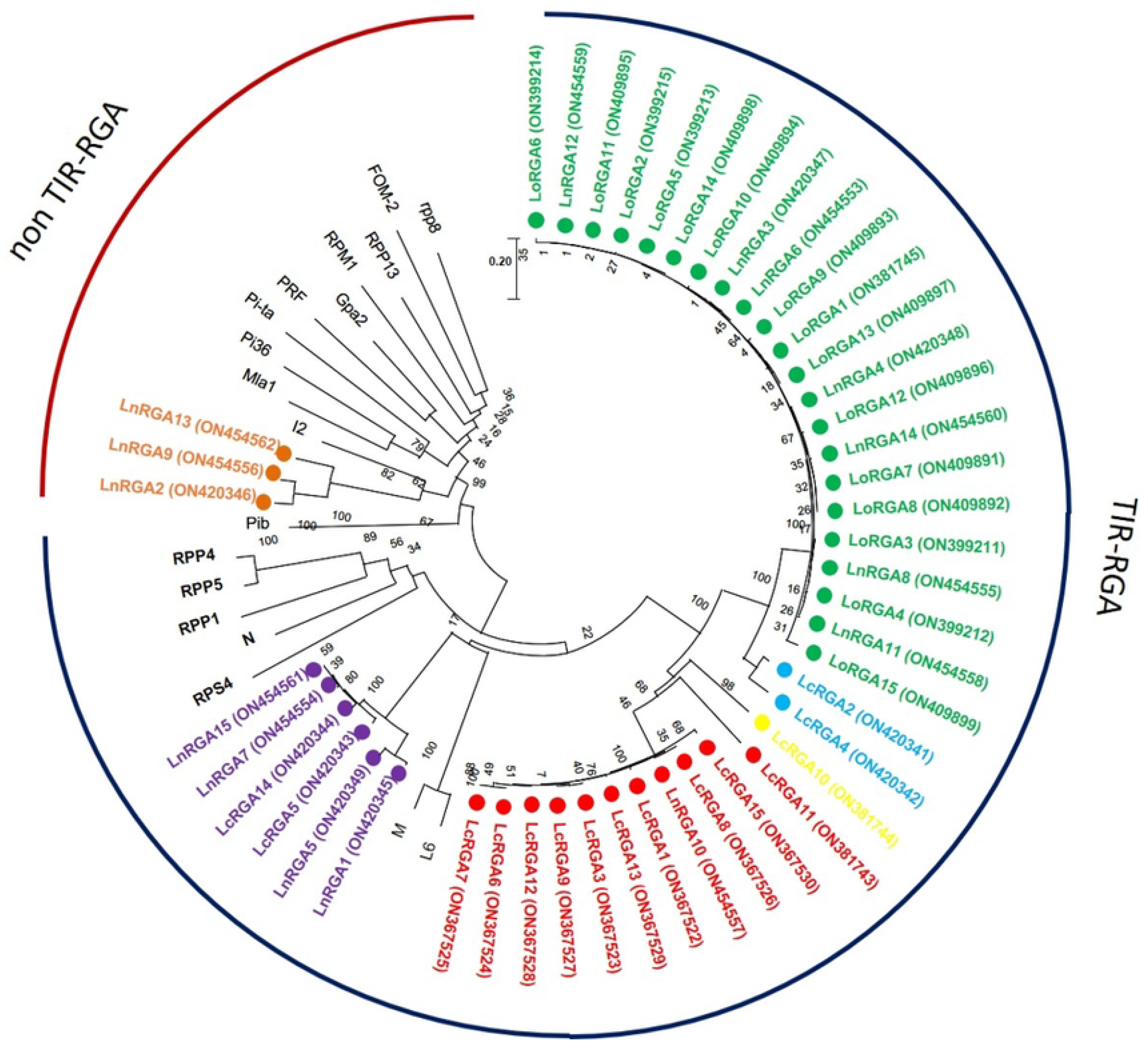
Neighbour joining phylogenetic tree constructed based on isolated Lentil RGAs and known NBS region of R gene, N (U15605), L6 (U27081), M (U73916), RPP5 (AAFO8790.1), RPP4 (AAM18462.1), RPP1 (AT3444670), RPS4 (CAB50708.1), Mla (AAG37356), Pi-ta (ACY24970.1), Pi36 (ADF29629.1), Pib (BAA76282.2), I2 (AF004878), RPP13 (AAF42831), RPM1 (AQ39214), Prf (U65391), Gpa2 (AF195939), RPP8 (AAC78631.1.), FOM-2 (AY583855.1) at bootstrap values (1000 replicates). Numbers on the branches indicate the percentage of bootstrap replications. Green represents LRGA 1, blue represents LRGA 2, yellow represents LRGA 3, red represents LRGA 4, purple represents LRGA 5, orange represents LRGA 6.

The Percent similarity of amino acid among lentil RGA and between R gene, L6, M, I2 and Gpa2 was determined using DNAMAN 8 software. Amino acid similarity ranged from 27.85% (LcRGA2 and LnRGA2) to 86.98% (LnRGA1 and LcRGA5) among RGAs. Similarity ranged from 26.83% (LnRGA13 and L6) to 49.41% (LnRGA13 and I2) when compared with the known R genes (Table 4).

**Table 4:** Homology matrix obtained between representative lentil RGA and known R gene using DNAMAN 8.0.

|  |  |  |  |  |  |  |  |  |  |  |  |  |  |
| --- | --- | --- | --- | --- | --- | --- | --- | --- | --- | --- | --- | --- | --- |
| LoRGA6 | 100 |  |  |  |  |  |  |  |  |  |  |  |  |
| LcRGA2 | 83.35 | 100 |  |  |  |  |  |  |  |  |  |  |  |
| LcRGA10 | 55.29 | 57.65 | 100 |  |  |  |  |  |  |  |  |  |  |
| LcRGA15 | 61.18 | 62.72 | 63.31 | 100 |  |  |  |  |  |  |  |  |  |
| LcRGA11 | 53.53 | 57.75 | 57.65 | 62.95 | 100 |  |  |  |  |  |  |  |  |
| LnRGA1 | 39.76 | 38.46 | 40.12 | 37.95 | 37.95 | 100 |  |  |  |  |  |  |  |
| LcRGA5 | 42.33 | 41.32 | 42.52 | 40.24 | 41.67 | 86.98 | 100 |  |  |  |  |  |  |
| LnRGA2 | 30.77 | 27.85 | 29.68 | 30.26 | 28.77 | 36.88 | 31.97 | 100 |  |  |  |  |  |
| LnRGA13 | 31.56 | 32.69 | 30.38 | 31.16 | 30.52 | 32.86 | 34.75 | 36.20 | 100 |  |  |  |  |
| I2 | 30.38 | 31.61 | 30.82 | 32.28 | 29.03 | 30.92 | 32.03 | 44.71 | 49.41 | 100 |  |  |  |
| L6 | 34.32 | 35.53 | 34.71 | 30.54 | 35.12 | 37.42 | 38.03 | 34.44 | 26.83 | 34.18 | 100 |  |  |
| M | 34.91 | 37.95 | 36.59 | 33.54 | 34.12 | 38.79 | 40.37 | 34.44 | 26.99 | 36.48 | 81.98 | 100 |  |
| Gpa2 | 27.81 | 32.69 | 28.13 | 31.29 | 30.41 | 32.86 | 34.75 | 36.20 | 42.14 | 37.87 | 33.33 | 32.03 | 100 |

### Motif identification and characterization

Motifs of TIR and non-TIR RGA were determined using Multiple Expectation maximizations for Motif Elicitation software along with known R gene. Eleven and twelve motifs were identified in TIR and non-TIR groups respectively. Six conserved motifs, P-Loop, RNBS-A, Kinase 2, Kinase 3, RNBS-C and GLPLA were found in all RGAs. External motif, P-Loop and GLPLA and internal motif, Kinase 2, Kinase 3 and RNBS-C were found conserved in both groups of RGA. We further classified RNBS-A to TIR with amino acid sequence [(YC)(AND)(RLK)I(SA) (NQDH)QF(EVDH)(AGM)(CSL)C(FL)(ILV)(DH)(DN)(NI)(SRG)] and non-TIR [F(CVD)(YL)(RK)(GAR)(WK)(SFA)(HTL)Y(SP)(KQE)(DVE)(YFL)(DC)(VA)(VRF)(TNA)(VI)] group due to the difference in position and composition of motif. The presence of Tryptophan (W) in the non-TIR group at the end of Kinase 2 motif distinguishes it from the TIR group with Aspartate (D) amino acid. An extra motif with signature (KE)NYRLH and E-value, 0E-054 was found in LcRGA 1, 3, 6, 7, 8, 9, 12, 13, 15 and LnRGA10 all belonging to class LRGA3 of TIR-RGA (Figs 6 and 7).

**Fig 6.**
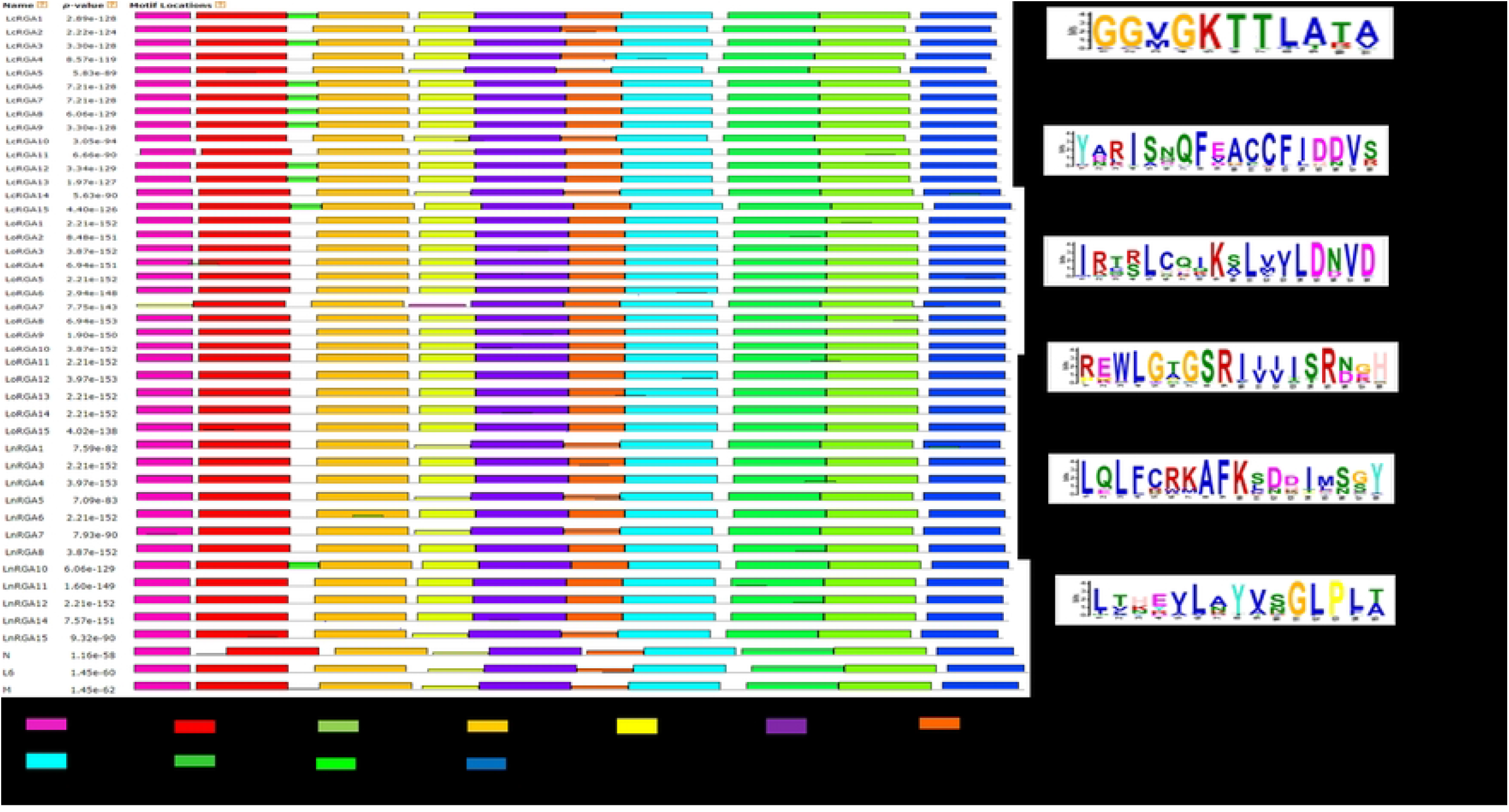
Diagrammatic representation of conserved motif of TIR RGAs within NBS domain along with R gene N, L6 and M. The solid black line represents each RGA and its length with motifs indicated in coloured boxes. The sequence logo of six conserved motifs along with their E value at right hand side.

**Fig 7.**
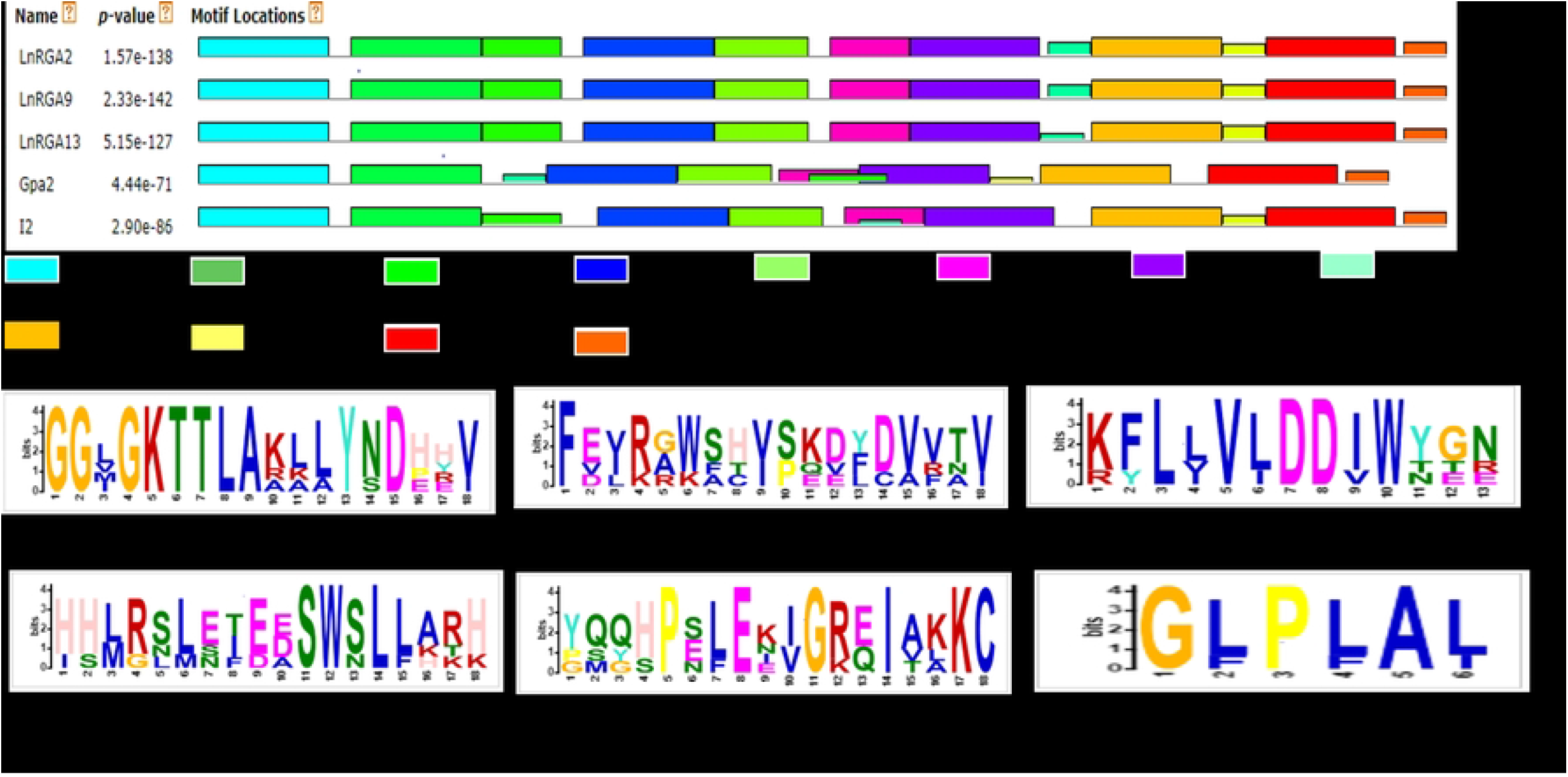
Diagrammatic representation of conserved motif of non-TIR RGAs within NBS domain along with R gene Gpa2 and I2. The solid black line represents each RGA and its length with motifs are represented in coloured boxes. Sequence logo of six conserved motifs along with their E value is given below.

### Prediction of active binding sites and their putative ligand

All the RGA showed structural analogy to know resistance gene, RPP1 and Roq1 with the molecular function of ATP binding (S4 Table). I-TASSER software was used to predict the active binding site of RGA and its respective ligand. The identified active binding sites (ABS) were found to be present along the six conserved motifs of Nucleotide binding site (NBS) including P-loop, RNBS-A, Kinas 2, Kinase 3, RNBS-C and GLPLA. ABS in P-loop motif was found in majorly 35 RGA (10 LcRGA, 14 LoRGA and 11 LnRGA) followed by Kinase 2 motif in 30 RGA (5 LcRGA, 14 LoRGA and 11 LnRGA). Amino acid corresponding to Active binding site, Glycine (G) and Threonine (T) in the P-loop and Aspartic acid (D) and Aspergine (N) in the Kinase 2 motifs were found common in TIR and non-TIR RGA but differed with amino acid at Kinase 2 motif. It was observed non-TIR had Aspartic acid as ABS while TIR-RGA had Aspartic acid and Aspergine as ABS. P-loop, Kinase 2 and Kinase 3 motifs are reported to have ATP/GTP binding site in nucleotide binding site of R gene. Interestingly Aspartic acid (D) found in TIR-RNBS motif was found to be active binding site of 17 TIR-RGA majorly of LoRGA isolated from *Lens culinaris* subsp. *odemensis*. LoRGA3, LnRGA8 and LnRGA14 had Leucine (L) found in RNBS-C and Glycine (G) amino acid found in GLPLA motif of non-TIR RGA, LnRGA9 and LnRGA13 as active binding sites. ADP/ATP was found to be a potential ligand of RGAs with the molecular function of ATP binding and ATPase activity (S5 Table). Previous reports of RNBS-A, RNBSC and GLPLA motif involved in ATP binding and hydrolysis has not been reported. Based on amino acid corresponding to ABS RGAs were classified into thirteen groups (Table 5). Eighteen RGAs with the active binding site at 4th, 6th, 27th and 82nd/83rd positions corresponding to G, T, D and N amino acid were grouped to GTDN class. Seven classes of ABS had a single lentil RGA describing the diversity of ABS within the genus. Seven RGAs had no ABS and were considered inactive due to point mutation. Little correlation was found between ABS of RGA and previously constructed phylogenetic tree. The RGA belonging to the non-TIR group had different ABS from TIR group. The phylogenetic tree was constructed by maximum likelihood in MEGAX aligning all the active binding sites of RGA with a bootstrap of 1000 replication. The tree grouped all RGAs having GTDN active binding sites together. RGA, LnRGA9, LnRGA13 (GTDT), LnRGA2 (GTDG) were grouped along with GTD group probably due to change in single amino acid (Fig 8).

**Table 5.**
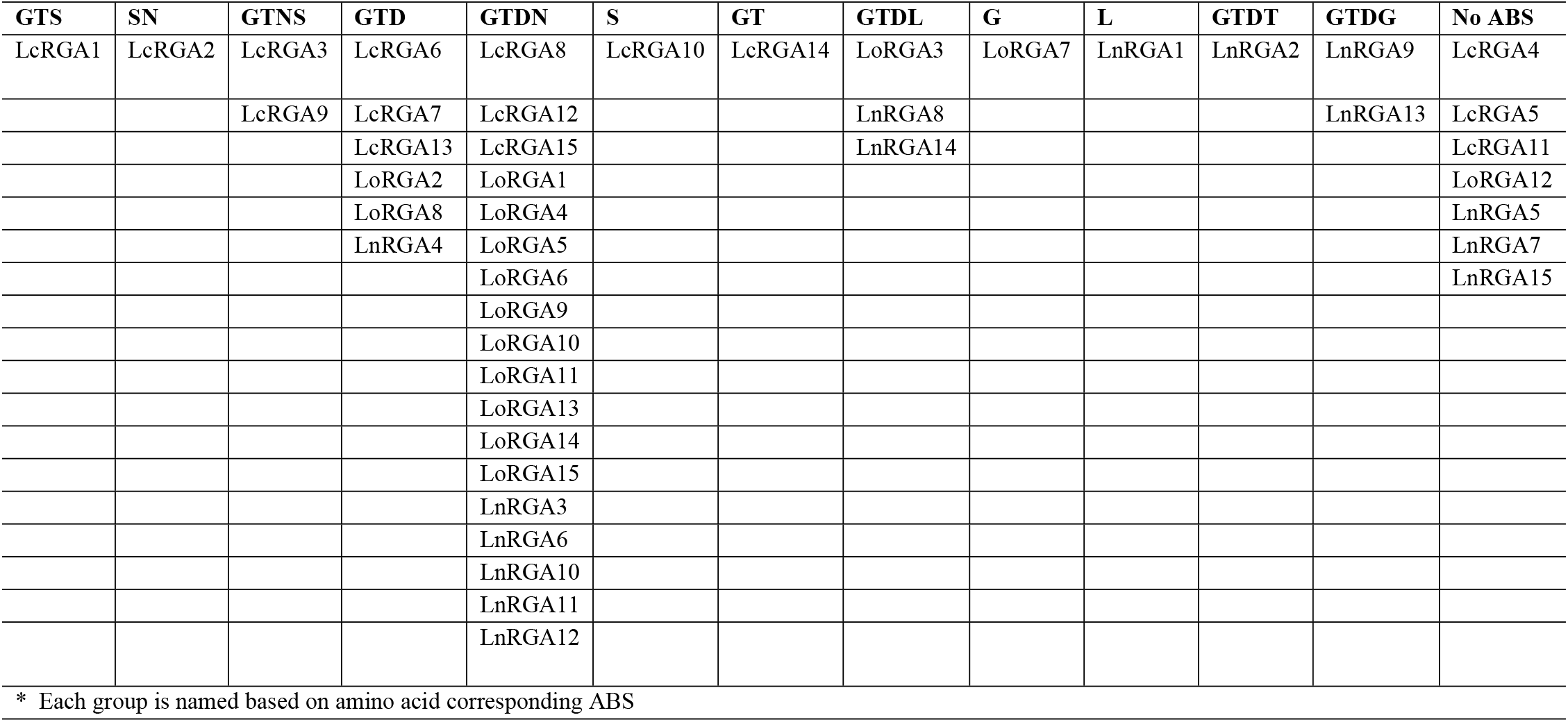
Lentil RGA grouped based on amino acid corresponding predicted active binding sites (ABS)

**Fig 8.**
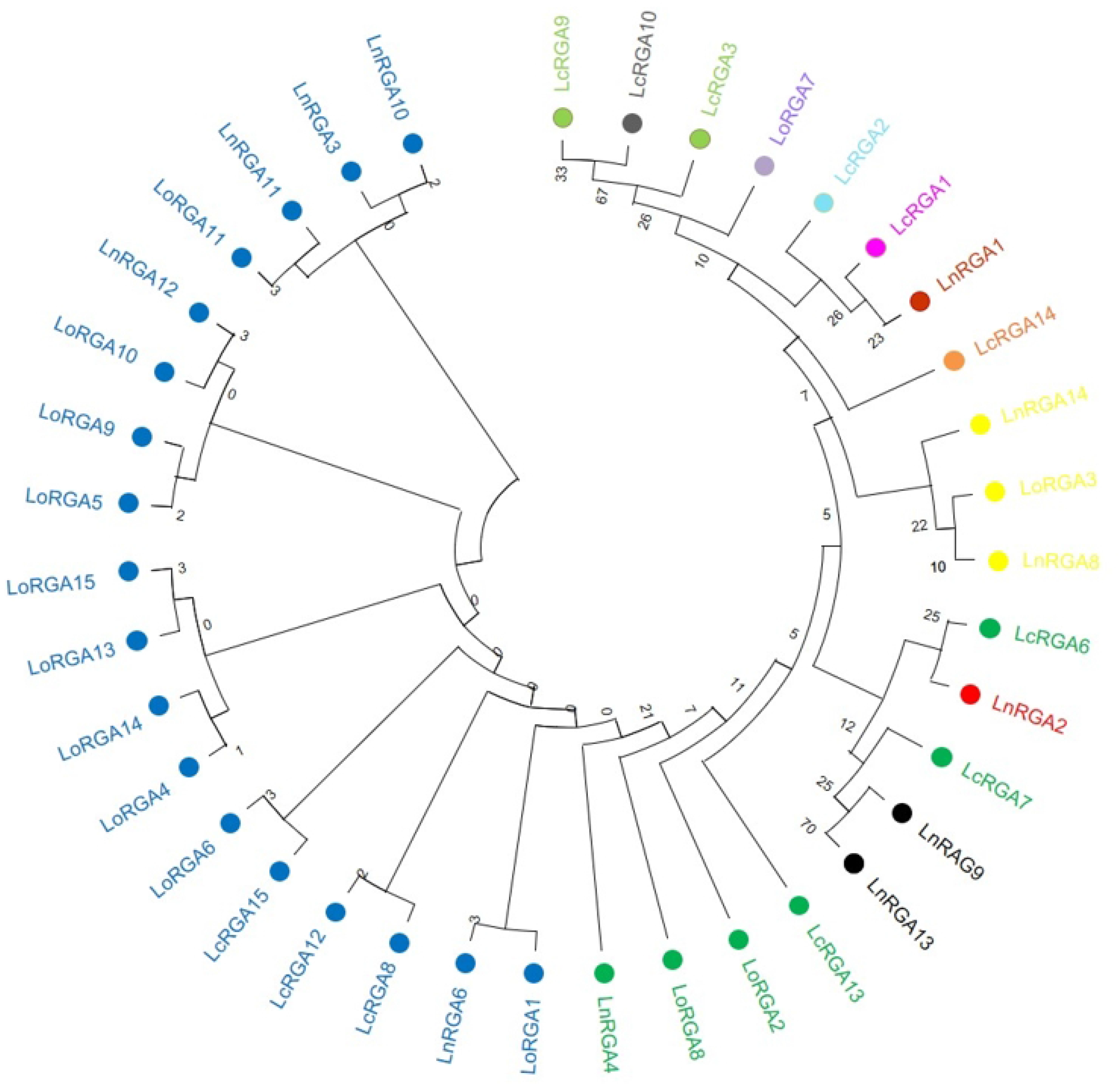
Phylogenetic tree constructed based on amino acid corresponding to active binding sites (ABS) using Maximum likehood in MEGA X with 1000 bootstrap replication. Lentil RGA with same colour represent having similar ABS. Blue represents GTDN, green represents GTD, black represents GTDG, Red represents GTDT, yellow represents GTDL, orange represents GT, red represents L, pink represents GTS, light blue represents SN, purple represents G, light green represents GTNS, grey represents S.

### The secondary and tertiary structure of RGA

The secondary structure of RGAs predicted using Pyre2 software revealed the presence of alpha-helix (56%-50%), beta strands (11%-9%), and disordered sequence (12%-14%). Three RGAs, LcRGA5, LcRGA11 and LnRGA1 had the transmembrane region in the helix depicts probable interaction with the lipid bilayer (Table 6). Best tertiary structure lentil RGA predicted based on C-score, TM-score and RMSD using I-TASSER. The C-score of 45 lentil RGA ranged from 0.31 (LcRGA14 and LoRGA2) to 0.68 (LnRGA9) and TM-score ranged from 0.70 ±0.10 (LnRGA11) to 0.81 ±0.09 (LnRGA9 and Lnrga13) (S6 Table). Solubility of active binding site amino acid ranged from 1-4 revealing the buried nature of ABS. It was predicted that RGA were isolated from genomic DNA represent constitutively inactive/closed conformation in nature until pathogen attack (S5 Figure).

**Table 6.** Secondary structure composition of representative lentil RGA using Pyre 2 software.

| Lentil RGA | $\alpha$ strand | $\beta$ strand | Disorder | TM helix | Confidence |
| --- | --- | --- | --- | --- | --- |
| LoRGA6 | 52 | 10 | 14 | - | 100 |
| LcRGA2 | 51 | 11 | 14 | - | 100 |
| LcRGA10 | 54 | 9 | 13 | - | 100 |
| LcRGA15 | 56 | 9 | 12 | - | 100 |
| Lcrga11 | 53 | 9 | 13 | 9 | 100 |
| LnRGA1 | 50 | 10 | 12 | 9 | 100 |
| LcRGA5 | 51 | 11 | 14 | 9 | 100 |
| LnRGA2 | 49 | 9 | 19 | - | 100 |
| LnRGA13 | 49 | 9 | 16 | - | 100 |

## Discussion

*Fol* is the major pathogen of lentil constraining its yield and productivity in India and worldwide. Identification of resistant germplasm through screening helps in the development of resistant cultivar and identification of R gene involved in resistance mechanism. In the present study, lentil germplasms were screened in natural conditions against seven race representatives of *Fol* for two seasons to identify resistant germplasm and further isolation and characterization of potential RGA. Since Fusarium is soil-borne pathogen, pot evaluation is considered efficient and accurate as it takes less space, provides uniform inoculum load and limits interaction with other soil-borne pathogens such as *Rhizoctonia bataticola* and *Sclerotium rolfsii* causing synergistic effect [27]. To increase the efficiency and reduce variation of screening the germplasms were screened for two consecutive seasons, 2020 and 2021 [28]. The germplasm showed typical wilt symptoms, yellowing, drooping to wilting of the plant followed by death. The germplasm showed varying degrees of resistance between the races in two seasons. Wild accession belonging to *Lens culinaris* sub sp. *odemensis* showed resistance to all the races of the pathogen. Multi-race resistance in has been identified in crops such as tomato against *Fusarium oxysporum* f. sp. *lycopersici* race 1, 2 and 3 [29] and in melon against *Fusarium oxysporum* f. sp. *melonis* race 0, 1 and 2 [30]. We have observed germplasms of *L. c* sub. sp. *culinaris* and *L. culinaris* sub sp. *orientalis* exhibited diverse reactions from highly resistant to susceptibility this might be probably due to heterogeneity in genome structure of germplasm within single species and differential interaction of resistant genes towards particular race. Our results were in accordance with previous reports of [31] differential resistance in the core set of *Phaseolus vulgaris* germplasm to races 1, 2 and 4 of *Fusarium*. The contrasting response showed by accessions of *L. lamottei* towards race 3, 5 and 7 emphasizes the probable combinatorial interaction of multiple R gene in resistance response. Few accessions showed variation in the resistance response as wilt is highly dependent on temperature [32]. Higher CV has been observed in wild species due to diverse disease reaction and small sample size. Extensive screening has explored the potentiality of all species and subspecies of lentil against existing races of *Fol* providing an excellent source for R gene isolation. To our best knowledge this is the first report of multi-race resistance in lentil against Fusarium wilt.

In last the decade PCR based approach for isolation of gene using degenerate primer has been identified as a valuable tool over conventional technique and have been used in crop plants to isolate resistant gene analogues closely associated with R gene. In our study accessions showing resistant response to multiple races were used for RGA isolation and characterization. Ninety clones were isolated and heterogeneity within the isolated RGA amplicons was observed. Similar results were also reported in radish [33]. Forty-five RGAs showed considerable sequence variation and similarity to RUN1and RPP13 disease resistance gene of legume model plant, *Medicago truncata* and TMV resistance gene, N of *Trifolium pratense* predicting its role in disease resistance. Our results were in accordance to RGA isolated from chickpea [34]. The presence of conserved domain P-loop, RNBS-A, Kinase 2, Kinase 3, RNBS C and GLPLA through multiple sequence alignment suggested it to be part of the NBS region of R gene. Amino-terminal of typical R gene is linked to TIR (Toll/interlukin 1-like receptor) and CC (Coiled-Coil) involved in defence signaling. They are differentiated based on the presence of aspartate or tryptophan (W) at end of kinase 2 motif in TIR and CC respectively [35]. We observed that forty-two isolated clones belonged to class TIR and the rest three to non-TIR. The presence of both classes of RGA and enhanced expression of TIR-NBS-LRR R gene have been reported in dicots and have evolved mainly through duplication and diversification during evolution [36-37]. In the present study, significant difference in the number of lentil TIR-RGA and non-TIR RGA clones was observed. To visualize the relatedness of isolated RGA and other known R gene phylogenetic tree was constructed based on amino acid sequence. The tree differentiated the clones to two groups, TIR and non-TIR and further into six classes, LRGA1-6. Clustering of all the RGA isolated from *L. culinaris* subsp. *odemensis* to class LRGA1 revealing sequence homology among the clones and could be due to tandem and segmental duplication within the sequence. Clustering of RGA from cultivated and wild into same classes reveals conserved nature of R genes. RGA isolated from three species were grouped to six classes revealing diversity of RGA present in the host. The diversity of RGA might be aided by recombination and sequence exchange resulting in haplotypic diversity [36]. Grouping of the LnRGA 13 along with the I2 Fusarium resistant gene and sharing 49% amino acid similarity predicts functional role in Fusarium wilt resistance [38]. All the conserved motifs were identified in both the classes of RGA when compared with R gene. Positional and sequence variation of RNBS-TIR and RNBS non-TIR motif have been also observed in Allium RGA resistant to Fusarium basal rot [39].

The presence of active binding sites determines the functionality of RGA in resistance. In-sillico characterization have identified six conserved motifs of NBS to harbor ABS and grouping based of ABS determines the diversity of RGA and probably different mode of action. Nucleotide binding site (NBS)/NB-ARC region of R gene belongs to STAND (Signal transduction ATPase with numerous domain) superfamily protein involved in immunity and apoptosis [40]. The biological function of all the isolated LRGA determined using I-TASSER inferred their involvement in immunity and involvement of LoRGA7 in intrinsic apoptotic signalling. TIR/CC-NBS-LRR requires maturation for recognition of effector molecules and is mediated by Hsp90, a ATP dependent chaperone and other co-chaperones. In closed and auto inhibited state, TIR/CC and LRR are present in close proximity and are folded back to NBS-ARC core along with ADP. Upon recognition of effector, conformational changes allow the exchange of ADP to ATP, resulting in open structure and activation of downstream defense [41]. Involvement of NBS region of I2 R gene in binding to ATP with P loop, Kinase 2 and Kinase 3 as ATP binding sites and subsequent hydrolysis of ATP by confirming relatedness of NBS to ATPase super family has been reported [42]. In the present study, ADP/ATP was found to be potential ligand of lentil RGA with ATP binding and ATPase as molecular function and infers its functional role in defense response. Potential solubility of ABS determined hydrophobic nature of amino acid. We predicted RGA secondary and tertiary structure using Phyre 2 and I-TASSER. The composition of Alpha-helix, beta strands and disordered region varied between RGA and similar results were found in RGA isolates from watermelon resistant to Fusarium wilt [43]. The presence of transmembrane helix deduced the role in lipid bilayer interaction [44].

## Conclusion

In the present study cultivated and wild lentil species resistant to multiple races of *Fol* were identified by extensive screening. Resistance gene analogues were isolated from resistant accession belonging to three different species using degenerate primer. The phylogenetic analysis grouped RGA to six classes determining the diversity of RGA present in the host. Clustering of cultivated wild species RGA together revealed conserved nature of R gene. Grouping of RGA isolated from *L. nigricans* with I2 reveals its potentiality in Fusarium wilt resistance. Molecular and biological function reveals ATP binding and ATP hydrolyzing activity of lentil RGA confirming their relatedness to the functional R gene. The isolated RGA can be useful marker associated with R gene and further expression analysis determine its activity during pathogen interaction and decipher the molecular mechanism involved in resistance.

## Acknowledgments

Authors acknowledge Dr. Kuldeep Tripathi of Division of Germplasm Evaluation, ICAR-NBPGR for providing lentil germplasm seeds and Dr. H. K. Dikshit, Division of Genetics for providing resistant and susceptible checks. The first author, N.K is grateful to CSIR for CSIR NET fellowship.

## Supporting information

**S1 Table. Primers used in the amplification of lentil resistance gene analogues**

**S2 Table. Screening of hundred lentil germplasm against seven-race representative *Fusarium oxysporum* f. sp. *lenti*l in the year 2020-21**.

**S3 Table. Screening of lentil germplasm against seven races *Fusarium oxysporum* f. sp. *lentil* in the year 2021-22**

**S4 Table. Prediction of structural analog, molecular, biological and cellular function of isolated Lentil RGA using I-TASSER**

**S5 Table. Predicted of Active binding site of RGA and its corresponding amino acid, its solubility and conserved motif**

**S6 Table. Predicted C-Score, TM-Score and RMSD of RGA tertiary structure**

**S1 Figure. Pot evaluation of lentil accessions, A: IC201561 (L65); B: EC714243 (L83) and C: EC718238 (L90) against seven races of *Fusarium oxysporum* f. sp. *lentis* and its respective control**. Pot in the left-hand corner is control followed by race 1(MP-2), race2(UP-9), race 3(RJ-8) race 4(DL-1) race 5(CG-5) race 6(UP-12) and race 7(BR-7) showing resistance reaction

**S2 Figure. Restriction digestion profile of plasmid having resistance gene analogue as insert (^≈^500 bp) isolated from *Lens culinaris* sub sp. *culinaris* (L65) digested with EcoR1 enzyme ran on 1.2% agarose gel**. Lane M is 1kb ladder and lanes 1 to 15 consist of RGA, LcRGA1 to LcRGA15.

**S3 Figure Restriction digestion profile of plasmid having resistance gene analogue as insert (^≈^500 bp) isolated from *Lens culinaris* sub sp. *odemensis* (L83) digested with EcoR1 enzyme ran on 1.2% agarose gel**. Lane M is1kb ladder and lanes 1 to 15 consist of RGA, LoRGA1 to LoRGA15.

**S4 Figure. Restriction digestion profile of plasmid having resistance gene analogue as insert (^≈^500 bp) isolated from *Lens nigricans* (L90) digested with EcoR1 enzyme ran on 1.2% agarose gel**. Lane M is 1kb ladder and lanes 1 to 15 consist of RGA, LnRGA1 to LnRGA15.

**S5 Figure. Tertiary structure of RGA predicted using online I-TASSER software**. RGA, LcRGA1-15 isolated from *Lens culinaris* subsp. *culinaris*, LoRGA1-15 isolated from *L. culinaris* subsp. *odemensis* and LnRGA1-15 isolated from *L. nigricans*.

## Reference

1. Arumuganathan K, Earle ED. Nuclear DNA content of some important plant species. Plant Mol. Biol. Rep 1991, 9: 208–218.

2. FAO. Production Crops. FAOSTAT Statistics Database. http://www.fao.org/faostat/en/Food; 2018.

3. Duran Y, Fratini R, Garcia P, De La Vega MP. An inter subspecific genetic map of Lens. Theor. Appl. Genet. 2004, 108:1265–1273.

4. Kumar S, Kumar J, Singh S, Ahmed S, Chaudhary G, Sarker A. Vascular wilt disease of lentil: a review. Journal of Lentil Research 2010, 4: 1–14.

5. Jiskani AM, Samo Y, Soomro MA, Leghari ZH, Gishkori ZGN, Bhutto SH, et al. A destructive disease of lentil: Fusarium wilt of lentil. Plant Arch. 2021, 21:2117–2127.

6. Hiremani NS, Dubey SC. Race profiling of Fusarium oxysporum f. sp. lentis causing wilt in lentil. Crop Prot. 2018, 108:23–30.

7. Khare MN. Diseases of lentil. In Lentils, Webb, C., Hawtin, G., Farnham Royal: England, UK, 1981. pp. 163–172

8. Agrawal SC, Singh K, Lal SS. Plant protection of lentil in India. In: Erskine W, Saxena MC, editors. Lentil in South Asia. Aleppo, Syria: ICARDA; 1993. pp. 147–165

9. Bayaa B, Erskine W, Singh M. Screening lentil for resistance to Fusarium wilt: methodology and sources of resistance. Euphytica. 1997, 98: 69–74.

10. Singh M, Kumar S, Basandrai AK, Basandrai D, Malhotra N, Saxena DR, et al. Evaluation and identification of wild lentil accessions for enhancing genetic gains of cultivated varieties. PloS one 2020, 15: e0229554.

11. Meena JK, Singh A, Dikshit HK, Mishra GP, Aski M, Srinivasa N, et al. Screening of Lentil (Lens culinaris Medikus sub sp. culinaris) Germplasm against Fusarium Wilt (Fusarium oxysporum f. sp. lentis). Int. J. Curr. Microbiol. App. Sci 2017, 6: 2533–2541.

12. Nair RA, Thomas G. Isolation, characterization and expression studies of resistance gene candidates (RGCs) from Zingiber spp. Theor. Appl. Genet. 2007, 116: 123–134.

13. Yaish MW, Saenz de Miera LE, Perez de La Vega M. Isolation of a family of resistance gene analogue sequences of the nucleotide binding site (NBS) type from Lens species. Genome 2004, 47: 650–659.

14. Palomino C, Satovic Z, Cubero JI, Torres AM. Identification and characterization of NBS– LRR class resistance gene analogs in faba bean (Vicia faba L.) and chickpea (Cicer arietinum L.). Genome. 2006, 49: 1227–1237.

15. Agbagwa IO, Patil PG, Das A, Soren KR, Singh IP, Chaturvedi SK, et al. Identification, Characterization, and Phylogenetic analysis of Pigeonpea (Cajanus cajan L. Millsp.) Resistance Gene Analogs using PCR cloning and in silico methods. Genetics and molecular research 2018, 17: 1–14.

16. He CY, Tian AG, Zhang JS, Zhang ZY, Gai JY, Chen SY. Isolation and characterization of a full-length resistance gene homolog from soybean. Theor. Appl. Genet. 2003, 106: 786–793.

17. Garzón LN, Oliveros OA, Rosen B, Ligarreto GA, Cook DR, Blair MW. Isolation and characterization of nucleotide-binding site resistance gene homologues in common bean (Phaseolus vulgaris). Phytopathology 2013, 103: 156–168.

18. Sekhwal MK, Li P, Lam I, Wang X, Cloutier S, You FM. Disease resistance gene analogs (RGAs) in plants. Int. J. Mol. Sci. 2015, 16: 19248–19290.

19. Bayaa B, Erskine WA. screening technique for resistance to vascular wilt in lentil. Arab J. Plant Prot 1990, 8: 30–33.

20. Murray MG, Thompson W. Rapid isolation of high molecular weight plant DNA. Nucleic acids research 1980, 8: 4321–4326.

21. Saitou N, Nei, M. The neighbor-joining method: A new method for reconstructing phylogenetic trees. Molecular Biology and Evolution 1987, 4, 406–425.

22. Kumar S, Stecher G, Li M, Knyaz C, Tamura, K. MEGA X: Molecular Evolutionary Genetics Analysis across computing platforms. Mol. Biol. Evol. 2018, 35: 1547–1549.

23. Bailey TL, Boden M, Buske FA, Frith M, Grant CE, Clementi L, et al. MEME SUITE: tools for motif discovery and searching, Nucleic Acids Research 2009, 37: 202–208.

24. Yang J, Yan R, Roy A, Xu D, Poisson J, Zhang Y. The I-TASSER Suite: Protein Structure and function prediction. Nat. Methods 2015, 12: 7–8.

25. Zhang Y. I-TASSER server for protein 3D structure prediction. BMC Bioinformatics 2008, 9: 40.

26. Kelley LA, Mezulis S, Yates CM, Wass MN, Sternberg MJ. The Phyre2 web portal for protein modeling, prediction and analysis. Nat. Protoc. 2015, 10: 845–858.

27. Chaudhary RG, Saxena DR, Dhar V, Singh RK, Namdev JK. Prevalence of wilt-root rot and their associated pathogens at reproductive phase in lentil. Arch. Phytopathol. PlantProt. 2010, 43: 996–100

28. Sharma M, Ghosh R, Tarafdar A, Rathore A, Chobe DR, Kumar AV, et al. Exploring the genetic cipher of chickpea (Cicer arietinum L.) through identification and multi-environment validation of resistant sources against Fusarium wilt (Fusarium oxysporum f. sp. ciceris). Front. sustain. food syst. 2019, 3: 78.

29. Reis A, de Britto Giordano L, Lopes CA, Boiteux LS. Novel sources of multiple resistance to three races of Fusarium oxysporum f. sp. lycopersici in Lycopersicon germplasm. Crop. Breed. Appl. Biotechnol. 2004, 4: 495–502.

30. Alvarez JM, González-Torres R, Mallor C, Gómez-Guillamón ML. Potential sources of resistance to Fusarium wilt and powdery mildew in melons. HortScience 2005, 40: 1657–1660.

31. Brick MA, Byrne PF, Schwartz HF, Ogg JB, Otto K, Fall AL, et al. Reaction to three races of Fusarium wilt in the Phaseolus vulgaris core collection. Crop Sci 2006, 46: 1245–1252.

32. Ahmed S, Akem C, Bayaa B, Erskine W. Integrating host resistance with planting date and fungicide seed treatment to manage Fusarium wilt and so increase lentil yields. Int. J. Pest Manag 2002, 48: 121–125. doi:10.1080/09670870110097690

33. Yu X, Kang DH, Choi SR, Ma Y, Lu L, Oh, S. et al. Isolation and characterization of fusarium wilt resistance gene analogs in radish. 3 Biotech 2018, 8: 1–9.

34. Huettel B, Santra D, Muehlbauer F, Kahl G. Resistance gene analogues of chickpea (Cicer arietinum L.): isolation, genetic mapping and association with a Fusarium resistance gene cluster. Theor. Appl. Genet. 2002, 105: 479–490.

35. Pan Q, Wendel J, Fluhr R. Divergent evolution of plant NBS-LRR resistance gene homologues in dicot and cereal genomes. J. Mol. Evol. 2000, 50: 203–213.

36. Joshi RK, Nayak S. Perspectives of genomic diversification and molecular recombination towards R-gene evolution in plants. Physiol Mol Biol Plants 2013, 19: 1–9.

37. Meyers B, Kozik A, Griego A, Kuang H, Michelmore R. Genome wide analysis of NBS-LRR-encoding genes in Arabidopsis. Plant Cell 2003, 15: 809–834.

38. Sun D, Hu Y, Zhang L, Mo Y, Xie J. Cloning and Analysis of Fusarium Wilt Resistance Gene Analogs in Goldfinger Banana. Mol. Plant Breed. 2010, 7: 1215–1222.

39. Rout E, Nanda S, Nayak S, Joshi RK. Molecular characterization of NBS encoding resistance genes and induction analysis of a putative candidate gene linked to Fusarium basal rot resistance in Allium sativum. Physiol. Mol. Plant Pathol. 2014, 85: 15–24.

40. Danot O, Marquenet E, Vidal-Ingigliardi D, Richet E. Wheel of life, wheel of death: a mechanistic insight into signaling by STAND proteins. Structure 2009, 17: 172–182.

41. Takken FL, Goverse A. How to build a pathogen detector: structural basis of NB-LRR function. Curr. Opin. Plant Biol. 2012, 15: 375–384.

42. Tameling WI, Elzinga SD, Darmin PS, Vossen JH, Takken FL, Haring MA, et al. The tomato R gene products I-2 and MI-1 are functional ATP binding proteins with ATPase activity. The Plant Cell 2002, 14: 2929–2939.

43. Reddy AC, Lavanya B, Tejaswi T, Rao ES, Reddy DC. Isolation and characterization of NBS-encoding disease resistance gene analogs in watermelon against fusarium wilt. Current Science 2019, 117: 617–626.

44. Zviling M, Kochva U, Arkin IT. How important are transmembrane helices of bitopic membrane proteins. Biochimica et Biophysica Acta (BBA)-Biomembranes, 2007, 1768: 387–392.

